# Data-Driven Flow-Map Models for Data-Efficient Discovery of Dynamics and Fast Uncertainty Quantification of Biological and Biochemical Systems

**DOI:** 10.1101/2022.02.19.481146

**Authors:** Georgios Makrygiorgos, Aaron J. Berliner, Fengzhe Shi, Douglas S. Clark, Adam P. Arkin, Ali Mesbah

## Abstract

Computational models are increasingly used to investigate and predict the complex dynamics of biological and biochemical systems. Nevertheless, governing equations of a biochemical system may not be (fully) known, which would necessitate learning the system dynamics directly from, often limited and noisy, observed data. On the other hand, when expensive models are available, systematic and efficient quantification of the effects of model uncertainties on quantities of interest can be an arduous task. This paper leverages the notion of flow-map (de)compositions to present a framework that can address both of these challenges via learning data-driven models useful for capturing the dynamical behavior of biochemical systems. Data-driven flow-map models seek to directly learn the integration operators of the governing differential equations in a black-box manner, irrespective of structure of the underlying equations. As such, they can serve as a flexible approach for deriving fast-to-evaluate surrogates for expensive computational models of system dynamics, or, alternatively, for reconstructing the long-term system dynamics via experimental observations. We present a data-efficient approach to data-driven flow-map modeling based on polynomial chaos Kriging. The approach is demonstrated for discovery of the dynamics of various benchmark systems and a co-culture bioreactor subject to external forcing, as well as for uncertainty quantification of a microbial electrosynthesis reactor. Such data-driven models and analyses of dynamical systems can be paramount in the design and optimization of bioprocesses and integrated biomanufacturing systems.

## 1 INTRODUCTION

Computational models have become indispensable tools for understanding the complex behavior of biological and biochemical systems towards design and optimization of bioprocesses and integrated biomanufacturing systems [1]. Recently, there has been a growing interest in data-driven methods for modeling the uncertain and nonlinear dynamics of biochemical systems, as these models constitute the cornerstone of various model-based analyses and decision-making tasks such as experiment design, hypothesis testing and parameter inference [2, 3, 4]. Data-driven modeling is especially useful when it is formidable to derive first-principles descriptions for systems whose complex behavior can span over multiple length- and time-scales. Data-driven models have shown promise for inferring the dynamics of cellular systems and metabolic networks (e.g., [5, 6]). Hybrid models (aka gray-box models) that combine physics-based models with data-driven descriptions of unknown or hard-to-model phenomena have also proven useful for describing the complex behavior of biochemical systems [7, 8, 9, 10].

In this work, we focus on *data-driven discovery* of dynamical systems, whereby the goal is to learn directly the governing equations from system observations. A class of data-driven discovery methods for unknown systems relies on basic assumptions about the structure of the underlying equations [11]. To this end, a popular technique is based on sparse identification from dictionaries of possible governing terms [12, 13], which has been shown to be particularly useful when limited system observations are available. On the other hand, nonparametric modeling approaches relax the necessity of using a library of candidate terms [14]. Another class of methods for data-driven reconstruction of dynamics is based on dynamic mode decomposition [15, 16], which approximates the eigenvalues and eigenvectors of the Koopman operator [17] that describes the dynamics of nonlinear systems.

Although inception of the field of nonlinear system identification dates back to few decades ago [18], the advent of machine learning, in particular deep learning, for characterizing complex input-output relationships has reinvigorated the interest in this area. Most notably, physics-informed neural networks [19] and dynamics reconstruction via neural networks under noisy data [20] have shown promise for data-driven modeling of nonlinear dynamical systems. Recently, Qin *et al*. [21, 22] proposed a deep learning-based approach for data-driven approximation of integration operator of differential equations from observations of state variables. The usefulness of this approach for discovery of dynamics of biological systems has been demonstrated on several benchmark problems in [23], mainly since it removes the necessity of assumptions about the dynamic model structure.

Data-driven discovery methods can also be used for model-based uncertainty quantification (UQ) applications that rely on expensive-to-evaluate computational models. Predictions of the behavior of biochemical systems are generally subject to various sources of uncertainty due to unknown model structure, parameters, and/or initial and boundary conditions. Systematic and accurate quantification of the effects of these uncertainties on predictions of quantities of interest (QoIs) is crucial when using models for decision-support tasks. This has spurred development of a plethora of set-based [24] and probabilistic [25, 26] methods for forward and inverse UQ problems (e.g., [27, 28, 29, 30, 31]). However, the most commonly used UQ methods rely on Monte Carlo sampling [32], which can be intractable for expensive computational models of biochemical systems, especially when models consist of a large number of differential equations and/or have a large number of uncertain inputs.

Surrogate modeling is being increasingly used to facilitate complex UQ analyses that would otherwise be computationally prohibitive. The key notion in surrogate modeling is to construct a data-driven mapping between inputs to a system and the QoIs in a non-intrusive manner, in which the “data generating process,” e.g., a high-fidelity model, is treated as a black-box to generate as few training samples as possible [33]. Such a data-driven representation can be used as a computationally efficient surrogate for expensive computational models in order to predict the QoIs as a function of inputs. A variety of surrogate modeling techniques such a generalized and sparse polynomial chaos [34, 35], Kriging [36] and deep learning [37] have been successfully applied to various biological and biochemical systems (e.g., [31, 38, 39, 40, 41]). Nonetheless, a critical challenge in the majority of these techniques arises from capturing the time-evolution of the QoIs in an efficient manner. The most common approach, known as *time-frozen* surrogate modeling [42, 43], for predicting the time-evolution of QoIs relies on constructing separate surrogate models for all time points at which the QoIs must be predicted. As such, the “time-frozen” approach can be an inflexible and inefficient way of surrogate modeling for dynamical systems, especially in dynamic UQ and decision-making problems that hinge on making predictions over an adaptive sequence of time instants.

In this paper, we leverage the notion of flow-map (de)composition, as also investigated in [21, 22], for data-efficient discovery of system dynamics from experimental observations or high-fidelity simulation data. Conceptually, a flow-map is an analytical operator that maps the current state and input of a system to a future state based on exact integration of model equations over some specified time step. Numerical integration schemes for ordinary differential equations in fact seek to numerically approximate flow-maps to compute the time-evolution of state variables as a function of input variables. Here, we propose to approximate flowmaps in a data-driven manner via non-intrusive surrogate modeling, such that the resulting *data-driven flowmap* is a surrogate for differential operators of the differential equations governing a dynamical system. Hence, data-driven flow-map models are able to discover system dynamics irrespective of the unknown structure of model equations. In addition, data-driven flow-map models can address the above-described challenge of “time-frozen” approaches to surrogate modeling via circumventing the need for construction of separate surrogate models at different time instants. This can be especially useful for fast UQ and optimization-based analyses of dynamical systems that hinge on repeated runs of expensive computational models over a sequence of time instants.

We demonstrate the usefulness of data-driven flowmaps for discovery of system dynamics from data, as well as for fast UQ applications based on expensive computational models. In this work, sparse polynomial chaos Kriging [44] is used for data-driven approximation of flow-maps owing to its data efficiency, ability to approximate complex mappings and ability to quantify the uncertainty of model predictions. The versatility of datadriven flow-maps is first demonstrated via the discovery of the transient behavior of benchmark problems and a co-culture bioreactor using noisy data. Subsequently, we show how data-driven flow-maps can speedup forward and inverse UQ analyses of a dynamic microbial electrosynthesis reactor, achieving up to a 100-fold gain in computational speed.

## 2 METHODS

In this section, we present the idea of flow-map (de)composition for dynamical nonlinear systems. This is followed by a discussion on the surrogate modeling technique and data generation strategies used in this work for learning data-driven flow-map models.

### 2.1 Flow-map Compositions

Consider a dynamical, time-invariant, nonlinear system described by

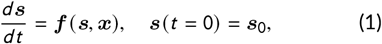

where 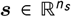 is the vector of state variables with initial conditions ***s***_0_, 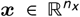 is the vector of input variables, and 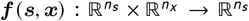 is the vector of (possibly unknown) system equations; ℝ denotes the set of real numbers. Eq. (1) describes the time-evolution of the state, ***s***, of the nonlinear system as a function of the inputs ***x***. Notice that in this work the inputs ***x*** can represent either model parameters, or manipulated input variables to a biochemical system, as will be discussed later.

A flow-map function is a mapping that predicts the transition of a dynamical system from the current to future state [21]. We define a flow-map function Φ_*δ*_ as

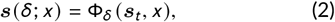

where ***s***_*t*_ denotes the current state at time *t* and *δ* is the lag time in the system transition from the current state ***s***_*t*_ to the future state ***s***. Given the current state of a system at time *t*, Φ_*δ*_ is in fact an analytical operation based on exact integration of ***f***, yielding the state after time *δ*

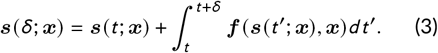

Eq. (3) describes the one-step transition between the states of a system. The integral term that appears in (3) can, subsequently, be considered as a flow-map residual, i.e., it represents the discrepancy between the current and future set of states.

The notion of flow-map compositions can be applied to compose a sequence of one-step transitions to define state trajectories over time [21]. Once the *δ*-lag flowmap Φ_*δ*_ is established, it can be used to predict states ***s*** at any time instant 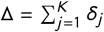 using a K-fold composition

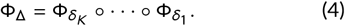

In practice, the set of differential equations in Eq. (1) describing the system dynamics may not be known, or, when known, their numerical solution may be expensive. In this paper, we aim to learn an approximate surrogate for the flow-map function in Eq. (2) from high-fidelity simulation or experimental data. Data-driven flow-map models can be established from simulation data to provide an efficient surrogate for expensive computational models of the form in Eq. (1) that, for example, rely on numerical integration of a large number of highly nonlinear and stiff differential equations, as is commonly the case for complex biochemical systems. Notice that in this case data-driven flow-map modeling can be viewed as approximating numerical time integrators of the differential equations in Eq. (1). Alternatively, in the absence of any knowledge about the governing equations (i.e., functions ***f*** in Eq. (1)), flow-map models can be directly learned from experimental observations in order to discover the unknown system dynamics. The main steps of data-driven flow-map modeling are summarized as follows. First, observations of the state variables are collected at several time instants either using highly-fidelity simulations, or via performing experiments. Notice that there is usually some degree of freedom in choosing the lag time *δ* in simulations, whereas the choice of *δ* is often limited by how fast measurements can be acquired in experiments. Then, the observations of the state trajectories over a sequence of discrete-time instants are used to train a surrogate for the flow-map in a non-intrusive, “black-box” manner. The data-driven flow-map model will take the states ***s***_*k*_, inputs ***x***_*k*_, and lag time *δ*_*k*_ at any discrete-time instant *k* as inputs to predict the future states ***s***_*k* +1_ at the time instant *k* + 1. With a slight abuse of notation, we denote the data-driven approximation of the flow-map in Eq. (2) by 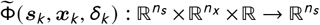. Figure 1 shows how a data-driven flow-map model can be used sequentially to predict the time-evolution of the states of a dynamical system. Notice that, at each time instant *k*, the flow-map model essentially “integrates” the states forward in time by *δ*_*k*_ until the final time is reached. Next, we discuss data-driven approximation of the flow-map.

**FIGURE 1.**
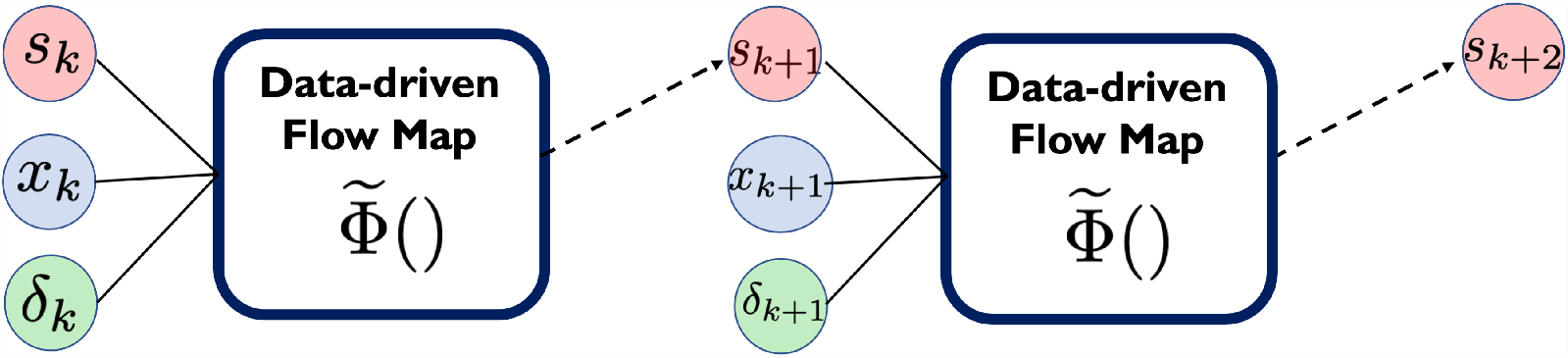
Data-driven flow-map models for predicting the state variables of a dynamical system over time. The flow-map model 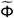 takes the current states ***s***_*k*_, inputs ***x***_*k*_, and lag time *δ*_*k*_ at a discrete-time instant *k* as inputs to predict the states ***s***_*k* +1_ at the subsequent time instant *k* + 1. By sequentially repeating this procedure, the time-evolution of the states in relation to the inputs can be established.

### 2.2 Data-driven Flow-maps

Here, we use sparse polynomial chaos Kriging (PCK) [44, 45] to learn a data-driven flow-map model 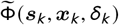 for the dynamical system in Eq. (1). Deep learning methods have also been used for approximating flow-maps for benchmark biological systems [23]. Yet, PCK combines the global approximation capability of polynomial chaos expansions, extensively used for surrogate modeling of (bio)chemical systems (e.g., [46, 47, 48]), with the local interpolation scheme of Kriging (i.e., Gaussian processes (GP) [49]). The polynomial structure of PCK makes its training data efficient, whereas Kriging offers the ability to quantify the uncertainties of model predictions.

Let us denote the vector of states, input variables, and lag time by 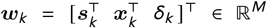, where *M* = *n*_*s*_ + *n*_*x*_ + 1. We represent ***w***_*k*_ as a multivariate random variable *W* with a (known) joint probability distribution *f*_*W*_, i.e., *W* ∼ *f*_*W*_. Notice that ***w***_*k*_ can be viewed as a realization of the random variable *W*; for notational convenience, we will drop the time index *k* in the remainder. The PCK approximation of the flow-map is defined as

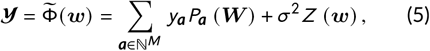

where 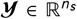 denotes the QoIs at *k* + 1 that are typically a subset of the states ***s***; *P*_***a***_ (***W***) are multivariate polynomial basis functions that are orthogonal with respect to the probability distribution *f*_*W*_ over the support 𝒟_*W*_ of the distribution, i.e.,

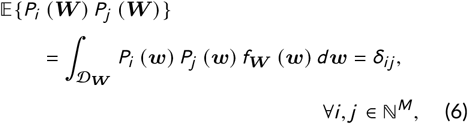

with 𝔼 being the expectation operator and *δ*_*ij*_ the Kronecker delta; *y*_***a***_ are the coefficients of the basis functions, with the multi-index ***a*** being an *M*-dimensional vector in the set of natural numbers ℕ; *Z* (***w***) is a standard normal random; and *σ*^2^ is a variance hyperparameter of the PCK.

The PCK in Eq. (5) represents the QoIs ***y*** as a Gaussian process (GP), such that the first term in Eq. (5) describes the trend (or mean) of the GP while the second term *Z* (***w***) describes the variance of the predicted QoIs. The trend of PCK is in fact an expansion of orthogonal polynomials that can represent any finite variance QoI [50]. Constructing the orthogonal basis *P*_***a***_ (***W***) requires the knowledge of the multivariate probability distribution *f*_***W***_. Eq. (6) gives the tensor product of *M* univariate polynomials that are orthonormal with respect to their corresponding marginal probability distribution. Optimal *L*_2_-convergence of the expansion of orthogonal polynomials has been established based on the Wiener-Askey scheme for various probability distributions [50, 51], although arbitrary orthogonal basis functions with sub-optimal convergence can also be constructed directly from moments of the random variable ***W*** [52]. As described, the multivariate random variable ***W*** consists of the states ***s***, input variables ***x***, and time lag *δ*. When ***x*** corresponds to uncertainties of a computational model (e.g., uncertainties in model parameters and/or initial conditions), their probability distribution is typically available *a priori* from parameter inference. As such, their respective polynomial basis functions can be chosen according to the Wiener-Askey scheme (e.g., Hermite basis for Gaussian distributions, Legendre for uniform distributions). On the other hand, when ***x*** corresponds to manipulated variables of a system, as is the case in the discovery of system dynamics, the input variables can typically be modeled as uniform distributions within a known range. The time lag *δ* can also be modeled as a uniform distribution within some range of interest for the application at hand. However, the distribution of states *s*_*k*_ is dependent on the realized state trajectories when the training data are generated and, thus, cannot be established *a priori*. Here, we assume states follow a multivariate Gaussian distribution with a mean and covariance computed from the training samples.

For practical reasons, the expansion of the trend term in Eq. (5) must be truncated up to a finite order. The truncated polynomial chaos expansion takes the form

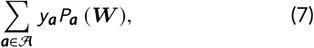

where the order of the expansion is dictated by the multi-index ***a*** ∈ 𝒜, with 𝒜 ⊂ ℕ ^*M*^ being the set of the multi-indices kept in the truncated expansion. The truncation scheme aims to limit the infinite expansion of the trend to a series of maximum order *p*. To address the challenges that arise due to increasing the order of the polynomial basis for better approximation and/or the large dimension of ***w***, sparsity can be introduced by employing the hyperbolic truncation scheme [35], also known as the q-norm scheme,

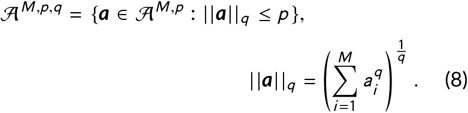

In principle, the coefficients *y*_***a***_ of the polynomial chaos expansion in Eq. (7) can be determined in a nonintrusive manner via solving a least-squares problem [53]. Here, we induce further sparsity by modifying the coefficient estimation problem to a *L*_1_-regularized regression problem [54]. The regularized coefficient estimation problem can be efficiently solved using the leastangle-regression (LAR) algorithm [55], which efficiently estimates the coefficients of the most relevant terms of the expansion in Eq. (7), setting the rest of the coefficients to zero.

Moreover, *Z* (***w***) in Eq. (5) is defined in terms of a kernel function *R* (|***w*** − ***w***′ |, *θ*), i.e., a function that provides some measure of similarity between different realizations of the random variable ***W***. Here, we use the Matérn kernel function [49]. Overall, the “tuning parameters” of the PCK that must be determined using the training data include the coefficients *y*_***a***_ of the trend, the variance term *σ*^2^, and the hyperparameters *θ* of the kernel function. This is efficiently done via maximumlikelihood estimation [44].

Finally, to quantify the quality of the PCK predictions, we use the leave-one-out cross-validation (LOOCV) error that is estimated from the training data. When onestep ahead test samples are available, validation errors can readily be evaluated. Furthermore, we assess the ability of the data-driven flow-map models in approximating the integration operator and, hence, their predictive accuracy over a multi-step integration horizon. Given *i* = 1, …, *N*_*V*_ validation state trajectories, each of which of length *T*_*i*_, we define the normalized, timeaveraged prediction error of QoIs, *ϵ*_*i*_, as

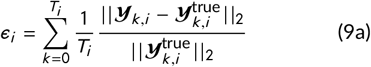

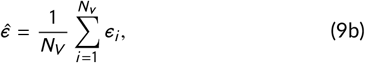

where | | · | |_2_ is the 2-norm of a vector; 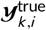 and ***y***_*k, i*_ are, respectively, the vector of OoIs in the validation dataset and those predicted by the data-driven flow-map models at time instant *k* for each validation run *i*. In the remainder, we refer to *ϵ*_*i*_ as the mean trajectory error (MTE), whereas 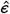 is the average MTE over all validation trajectories.

### 2.3 Data Generation and Model Training

To train an approximate flow-map model 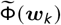, we require input-output data that represent one-step transitions between states. To this end, a total of *N*_*T*_ trajectories of state variables ***s***_*k*_ over a discrete-time horizon {0, 1, · · ·, *k, k* + 1, · · ·, *T*} are generated, where *T* is the length of the time horizon of the training trajectories. At each time instant *k*, a single training sample consists of ***w***_*k*_ → ***y***_*k*_.

For trajectory generation, it is crucial to vary the initial conditions ***s***_0_ and inputs ***x***_***k***_ within some allowable range, as well as the time lag *δ* whenever applicable. The training data must cover a wide range of state, input and time lag values, as relevant to the application of the trained models. As such, each sample of observed states within each trajectory represents a unique transition from the current to future state of the system for the given input and time lag values. We note that an effective strategy for generating simulation data is via one-step transitions. That is, instead of generating an entire trajectory given some initial conditions ***s***_0_, we can randomly sample the state-space, along with the uncertain parameters and time lag, in order to compute the corresponding future states.

The data generation and PCK model training strategy adopted in this work is summarized in Figure 2. We remark that, although random sampling is used here to generate the training data, PCK provides confidence estimates on its predictions that can be used towards active learning-based sampling (e.g., see [56]). As will be demonstrated in the subsequent sections, the main benefits of using PCK for constructing data-driven flowmap models include: (i) being more data efficient, especially as compared to feedforward neural networks [23], when used for discovery of system dynamics from system observations; (ii) offering significant improvements in the computational efficiency of data generation for surrogate modeling for dynamical systems as compared to time-frozen approaches; and (iii) characterizing the uncertainty of model predictions.

**FIGURE 2.**
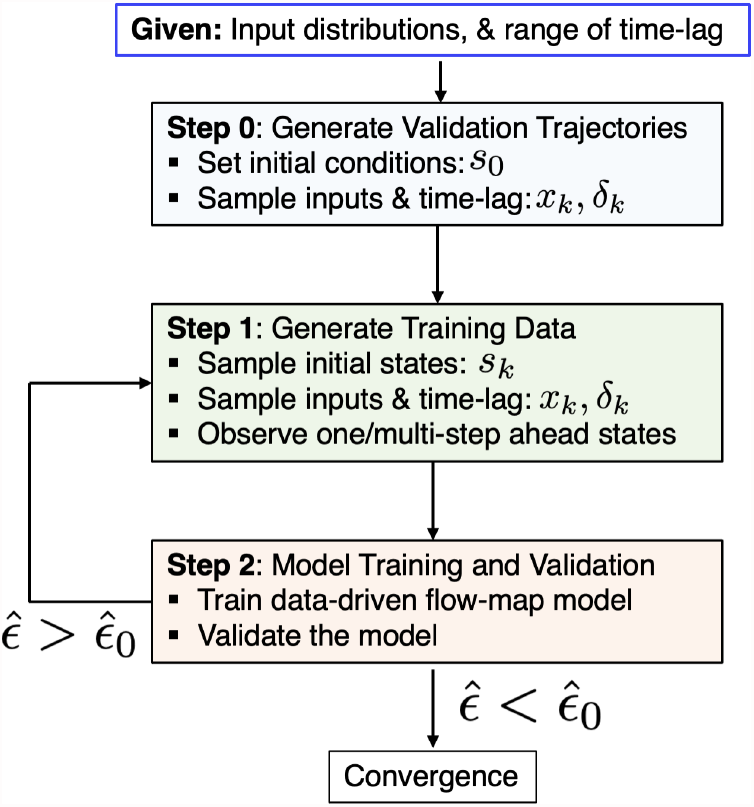
Algorithm for data generation and training of data-driven flow-map models. Validation trajectories are first generated. Then, one/multi-step ahead simulations or experiments are performed to observe successor states given the initial states, inputs, and time-lag. Subsequently, the data-driven flow-map model is trained. In the case of PCK models used in this work, several hyperparameters must be selected during the model training. These include the polynomial order, hyperbolic truncation parameter, covariance function and the regression method used for estimating the expansion coefficients. Finally, the prediction accuracy of the trained model is assessed against the long-time validation trajectories. If the prediction accuracy 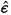 is larger than some pre-specified threshold 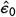, the model training and validation process will be repeated.

In this work, the following procedure is used for fitting the PCK models. We use the sequential PC-Kriging approach proposed in [44], where a polynomial chaos expansion (PCE) is first trained based on the available data and is then embedded as the trend of PCK. For training the PCE, we allow the polynomial expansion’s maximum order to vary from 1 to 5; higher order polynomials are avoided to retain a smaller expansion (i.e., less degrees of freedom) and mitigate overfitting. The truncation factor *q* in Eq. (8) is varied from 0.7 to 0.85 since the resulting maximum order of the polynomials will ensure that we do not have highly nonlinear interaction terms while allowing for elimination of few of interaction terms. The optimal value of *q* is chosen based on cross-validation. We use a Matérn kernel for the GP part of PCK models. The hyperparameters of PCK are selected using a data-driven optimization algorithm, namely the covariance matrix adaptation–evolution strategy [57].

## 3 DATA-DRIVEN DISCOVERY OF DYNAMICAL SYSTEMS

In this section, we apply the PCK-based flow-map modeling approach to learn the dynamics of several benchmark systems using limited data. The first case study, based on the Morris-Lecar system, compares the performance of the PCK model with neural network modeling results of [23]. The second case study, based on the Lorenz system, focuses on reconstructing the dynamics of a chaotic system in which variations in parameters significantly change the solution landscape. Lastly, we show how the flow-map modeling approach can be used for discovering the dynamics of a co-culture bioreactor under noisy observations and how the variance term of PCK provides a measure of uncertainty of model predictions.

### 3.1 Morris-Lecar System

The first benchmark problem is the Morris-Lecar system [58], which describes neuronal excitability. This system was used in [23] to examine neural network-based flowmap models for the discovery of nonlinear dynamics. In particular, a residual neural network was used to represent the data-driven flow-map model, in which only the flow-map residual is learned by skipping the input connection to the neural network and adding it to the output of the latter. Here, we aim to recreate the results of the aforementioned work, demonstrating the data efficiency of the proposed PCK approach to data-driven reconstruction of dynamics. The dynamics of the Morris-Lecar system are described by

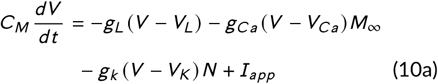

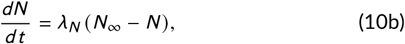

where *V* (mV) is the voltage difference between the sides of the membrane and *N* represents the probability for the potassium channel being open. The parameters *M*_∞_, *N*_∞_ and *λ*_*N*_ depend on the voltage, as defined in the SI. We focus on the so-called Type I model with parameters taken from [23] and given in the SI. Here, it is assumed that the model parameters are fixed, as we aim to reconstruct the system dynamics as a function of *x*_*k*_ = *I*_*app*_ that can vary within the range [0, 300] A. Specifically, we aim to predict the long-term system dynamics, starting from a given initial conditions, under a fixed *I*_*app*_. To compare our results with those in [23], *δ*_*k*_ was chosen to be 0.2 ms; we did not consider the time-lag as part of the PCK model. This system exhibits a saddle node bifurcation, which leads to an oscillatory behavior depending on the value of input *I*_*app*_. Thus, the data-driven flow-map model must capture the oscillatory behavior for different values of *I*_*app*_.

To train the PCK-based flow-map model, we generated one-step ahead samples of the states *V*_*k*_ and *N*_*k*_ by randomly drawing the initial states from [−75, 75] × [0, 1]. Here, we first examine the convergence error of the flow-map model to characterize how many samples of states would be necessary for data-driven reconstruction of the system dynamics. We quantify the convergence error in terms of the average MTE in Eq. (9) based on three validation trajectories generated for *I*_*app*_ = {0, 60, 150}. Figure 3 shows the average MTE estimated over 1,000 time steps in relation to the number of training samples, where the vertical line around each error represents one standard deviation based on 5 repetitions of the analysis. It is evident that the error converges after about 160 samples, suggesting that a limited number of training samples is needed.

**FIGURE 3.**
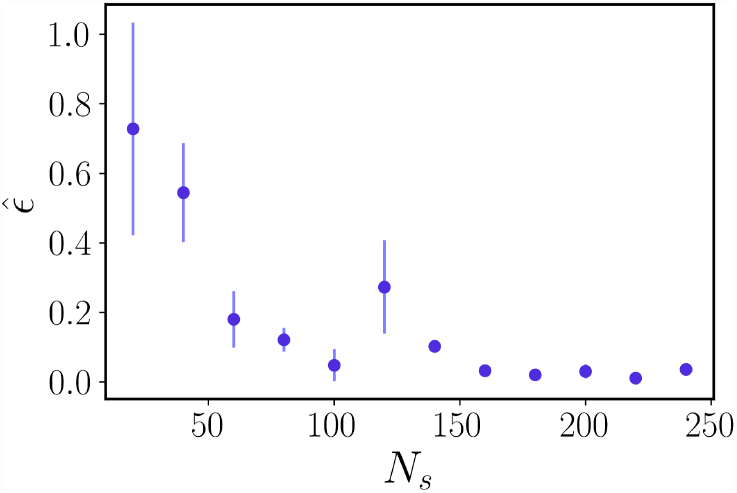
The average mean trajectory error, 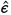, of the PCK-based flow-map model for the Morris-Lecar system in relation to the number of training samples, *N*_*s*_. The error is estimated based on three validation trajectories generated for the input *I*_*app*_ values {0, 60, 150}. The vertical bars represent the standard deviation of the error estimated based on 5 repeats of the training.

Figure 4 shows the reconstructed dynamics by the PCK-based flow-map model trained using 240 samples in comparison with the true dynamics. As can be seen, there is no visible discrepancy between the true time-evolution of the system and the reconstructed dynamics. The system exhibits a bifurcation behavior, as evident from the phase plots shown in Figure 4(c), (f), (i). Yet, the PCK-based flow-map model is able to capture this complex behavior and accurately predict the system dynamics over a long-time horizon. We note that a 500-fold saving in the number of training samples is observed as compared to [23] in which a recurrent neural network representation was used for the flow-map model. This is while the PCK model also yields slightly more accurate predictions.

**FIGURE 4.**
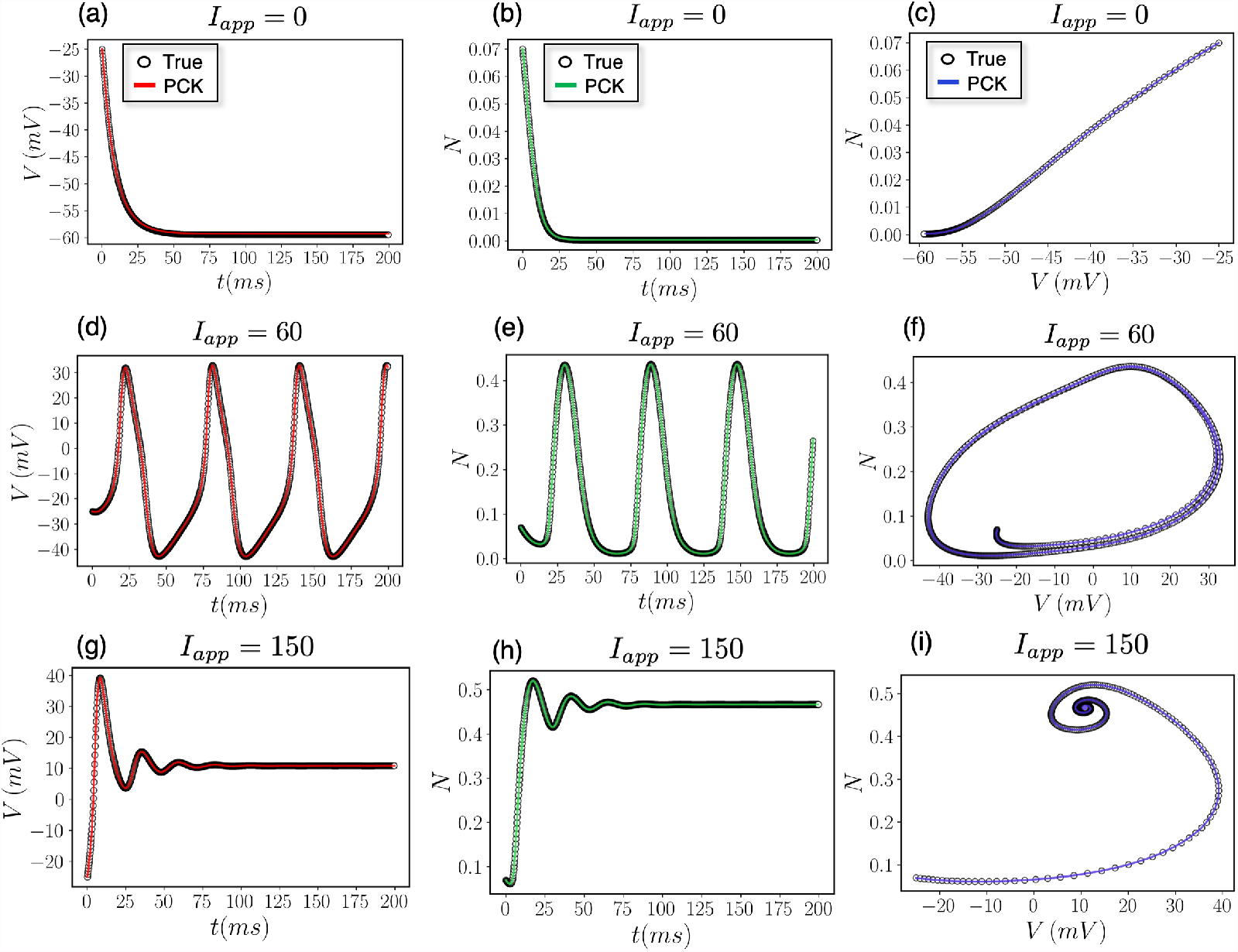
Reconstructed dynamics of the Morris-Lecar system by the PCK-based flow-map model in comparison with the true system dynamics for the input *I*_*app*_ values {0, 60, 150}. The PCK-based flow-map model is trained using 240 samples. The left column shows the time-evolution of voltage difference, *V*; the middle column shows the time-evolution of the channel opening probability, *N*; and the right column shows the corresponding phase plots.

### 3.2 Lorenz System

We now consider a chaotic dynamical system based on the well-known Lorenz benchmark problem [59]. The Lorenz system has been widely used in the data-driven modeling literature (e.g., [60, 61]). The Lorenz system is described by the following set of nonlinear ordinary differential equations

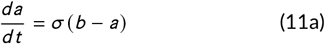

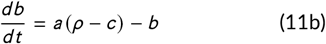

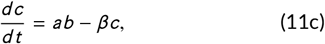

where ***s*** = [*a, b, c*]^⊤^ are the system states and ***x*** = [*σ, ρ, β*]^⊤^ are the uncertain model parameters. Chaotic behaviors can be encountered in various chemical and biological systems, including in the growth of biological populations with non-overlapping generations [62] and the peroxidase–oxidase oscillator [63]. Here, we consider a constant time-lag *δ* = 0.01 that captures the intrinsic time-scale of the system [64].

The Lorenz system exhibits a chaotic behavior based on the initial conditions ***s***_0_, while its long-term behavior is highly affected by the uncertain parameters ***x***. The nominal initial conditions and parameters of the system are, respectively, ***s***_0_ = [1.9427, −1.4045, 0.9684]^⊤^ and ***x***_0_ = [10, 28, 8/3]^⊤^, for which the system oscillates around two attractors. Here, the training data consisted of 500 random samples of the state-space ***s*** within the range [−10, 10] × [−10, 10] × [−10, 10] and the parameters ***x*** within the range [8, 12] × [10, 30] × [1, 5.5]. We used two validation trajectories to compare the true system dynamics with those reconstructed by the PCK-based flow-map model: one trajectory based on the nominal initial conditions and parameters and the other based on ***x*** = [10, 15, 8/3]^⊤^ and ***s***_0_ = [1.6655, −0.1178, 0.1748]^⊤^.

Figure 5 shows phase plots of the reconstructed oscillatory dynamics of the Lorenz system, in comparison with the true system dynamics, over a simulation horizon of 5,000 time steps. We observe that the qualitative behavior of the Lorenz system is different when the parameter *ρ* is varied, while the PCK-based flowmap model is able to reconstruct the dynamics in both cases. The MTE is 0.522 for the nominal validation trajectory and 0.0013 for the second validation trajectory. Although the error for the nominal validation trajectory seems relatively high, the main characteristics of the true dynamics are adequately captured, as evident from Figure 5(a)-(c). That is, the limit circles, the amplitude of oscillation and period are adequately captured. These predictions are consistent with those reported in [61]. However, we note that reconstruction of the Lorenz dynamics using neural networks typically requires on the order of a few thousands of training samples [20, 64], whereas the PCK model here was trained using 500 samples.

**FIGURE 5.**
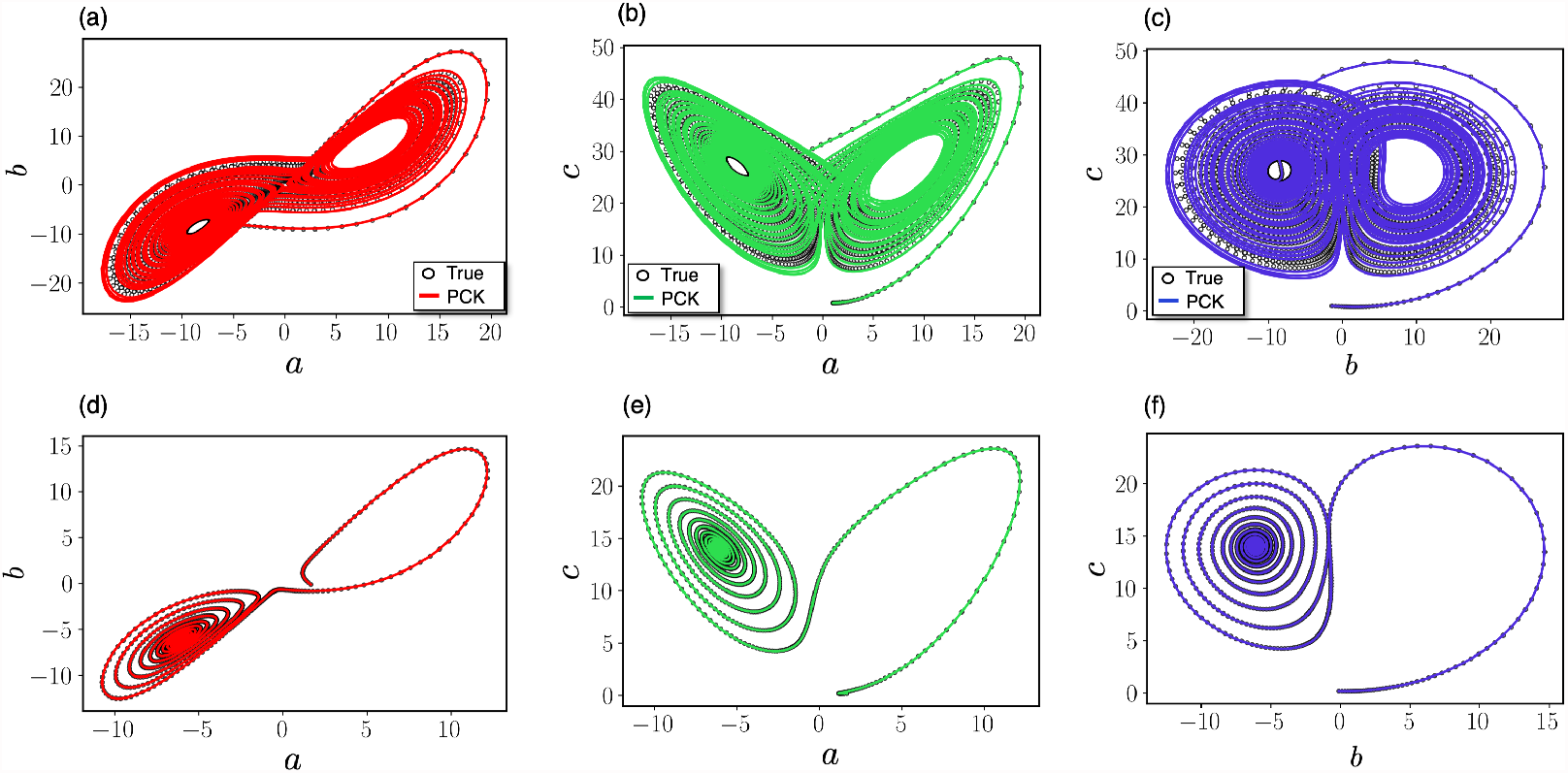
Phase plots of the reconstructed dynamics of the Lorenz system by the PCK-based flow-map model in comparison with the true system dynamics for different values of model parameters. Subplots (a)-(c) correspond to the model parameters *σ* = 10, *β* = 8/3, and *ρ* = 28. Subplots (d)-(f) correspond to the model parameters *σ* = 10, *β* = 8/3, and *ρ* = 15.

### 3.3 Transient Co-culture System

In this case study, we demonstrate the ability of PCK- based flow-map models to learn the transient behavior of a co-culture system with variable inputs. In particular, we focus on the startup dynamics of a continuous bioreactor driven by the competition of several auxotrophs [65]. To emulate data collection from a real system, we use a nonlinear dynamic model of the bioreactor [66] (given in the SI) to generate observations of the system states, which are then corrupted with independent and identically distributed state-dependent measurement noise 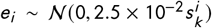, with *i* being an index for the measured states and *k* the time index. The five state variables ***s***_*k*_ of the bioreactor include: the population of the two species *N*_1_ (*Cel l s* /*L*) and *N*_2_ (*Cel l s* /*L*), the auxotrophic nutrients concentrations *C*_1_ (*g* /*L*) and *C*_2_ (*g* /*L*), and the common shared carbon source concentration *C*_0_ (*g* /*L*). The bioreactor has three process inputs ***x***_*k*_ that can be varied in time. The process inputs are the dilution rate *D* (hr^−1^) that varies within the range [0.75, 1.5] (hr^−1^), as well as the feed substrate concentration of auxotrophs *C*_1,*in*_ (g/l) and *C*_2,*in*_, both varying in the range [1.5, 2] (g/l). To generate data for training the PCK-based flow map models, short simulation “experiments” with a fixed length of *T* = 30 steps with *δ*_*k*_ ∈ [0.15, 0.25] hr^−1^ were performed. At each time step *k* during the multi-step experiments, inputs ***x***_*k*_ were varied over the time interval *δ*_*k*_ and noisy observations of the states were collected. For the validation plots of Figure 6, we begin by some random initial condition at *k* = 0, by applying an input *x*_0_ over some interval *δ*_0_. The model predicts the mean of the states at *k* = 1, as well as their variance. The integration proceeds by taking a next step based on the mean value of the states at *k* = 1, predicting the states at *k* = 2. Using only the mean value to compute trajectories is probably the simplest way when Gaussian Process state space models are utilized, however, there are more sophisticated ways for the trajectory generation [67], which are beyond the scope of the paper. Note that properly incorporating the uncertainty in multi-step ahead predictions is a complicated issue addressed in the literature [68, 69]. Here, it suffices to use a deterministic function, e.g., the mean value of the data-driven flow-map model, to integrate in time since this way we avoid the major issue of using noisy inputs into our PCK model. The validation trajectories have a length of *N*_*k*_ = 40 steps ahead, extending slightly beyond the training range. Moreover, thanks to the nature of the PCK model, we can also simply characterize the confidence of the model to the prediction of the dynamics. To get some uncertainty estimates on the predicted trajectories, at each step *k*, we plot the 3*σ* (*w*_*k*_) error bars around the mean. Overall, we observe that the true, noiseless trajectories are embedded within the confidence intervals of the PCK predictions.

**FIGURE 6.**
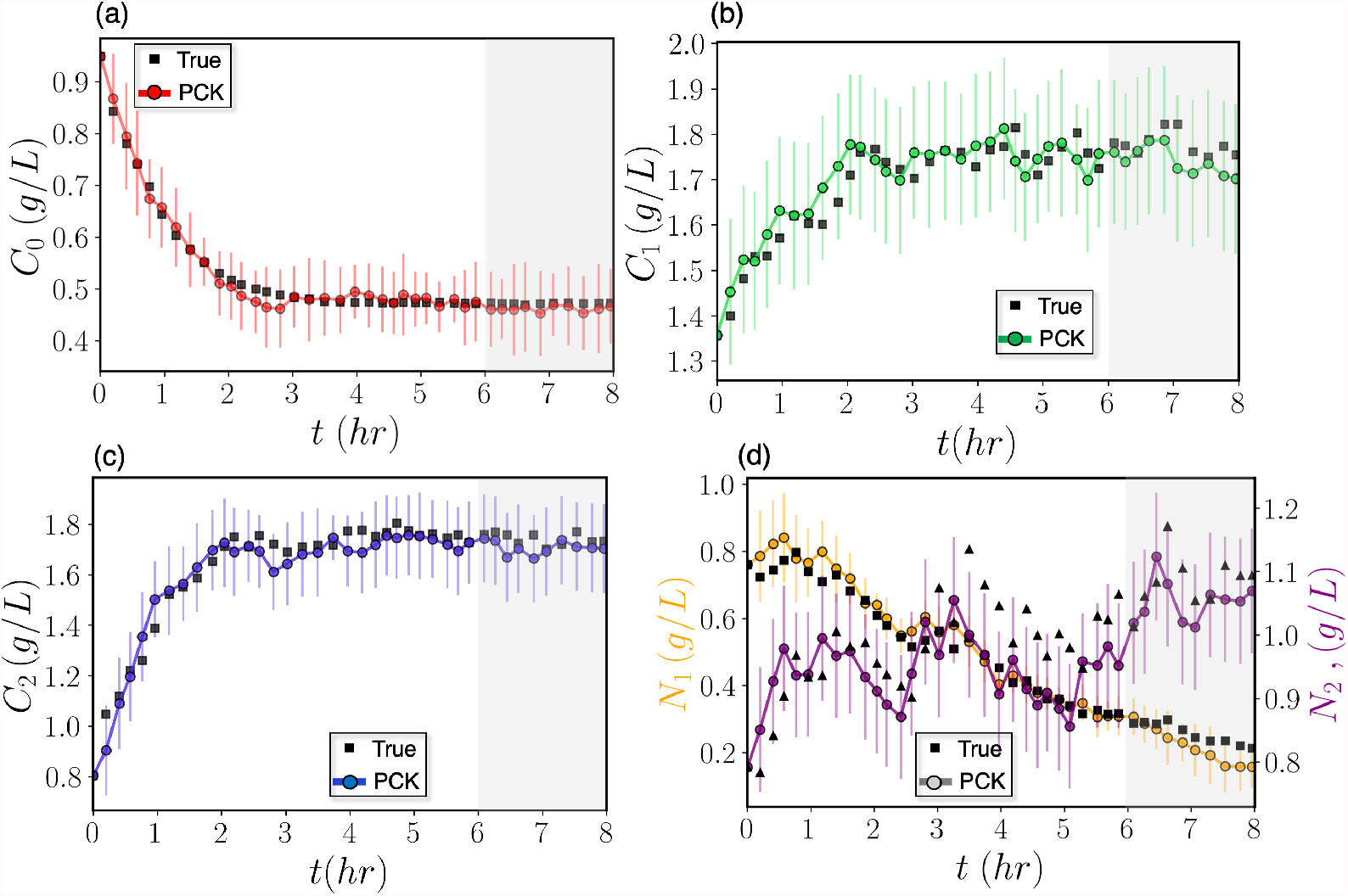
Predictions of the state variables of the transient co-culture system via the PCK-based flow-map models in comparison with the observed state trajectories. The colored lines/points correspond to the predicted trajectories by the mean of the PCK models, starting from some initial states at *t* = 0 hr. Black symbols represent the observed trajectories at specific snapshots during a validation run. Vertical error bars represent the uncertainty in the predictions of the PCK models, estimated as plus/minus two standard deviations from the mean value. The shaded areas correspond to a time interval that was not accounted for when training the PCK models.

## 4 UNCERTAINTY QUANTIFICATION OF EXPENSIVE COMPUTATIONAL MODELS

In this section, we demonstrate the utility of data-driven flow-maps for the UQ of a Microbial Electrosynthesis (MES) bioreactor using a high-fidelity computational model that is subject to uncertainty in model parameters and initial conditions. In particular, we show how flow-maps can be used as surrogate models for efficient sample-based approximation of distribution of QoIs, global sensitivity analysis, and Bayesian parameter inference, when the original model is prohibitively expensive for a sample-based analysis.

We consider the batch MES bioreactor shown in Figure 7 for CO_2_ fixation [70], with potential applications in space biomanufacturing [71]. The bioreactor consists of a well-mixed liquid bulk phase that contains dissolved CO_2_, i.e., substrate. A microbial community forming a biofilm grows on the cathode of the bioreactor. The dissolved substrate diffuses into the biofilm through a linear boundary layer and is then consumed by bacteria towards the growth of the biofilm. This leads to spatial distribution of the substrate concentration within the biofilm. Voltage is applied to the cathode while the biofilm acts as a conductive matrix through which electron transport takes place. Both the substrate CO_2_ in the biofilm and the local overpotential due to the current flux contribute to the biofilm growth kinetics described by the dual Monod-Nerst model [72].

**FIGURE 7.**
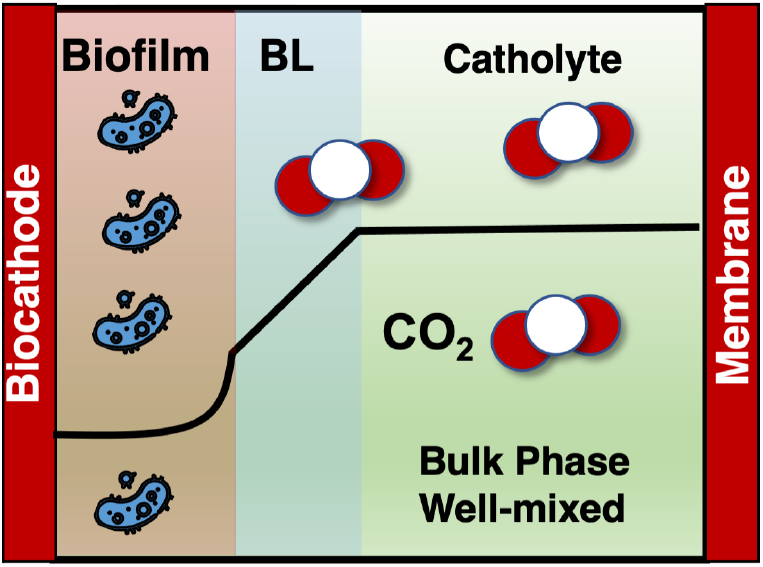
Schematic of the microbial electrosynthesis bioreactor. The bioreactor consists of 3 regions: the bulk phase, the biofilm, and a boundary layer (BL) in between. The black line represents a typical concentration profile of some species as predicted by the computational model used in this work. The concentration is assumed to be constant in the bulk phase, changing linearly across the boundary layer, and exhibiting a more complicated shape in the biofilm.

A computational model of the dynamics of the MES bioreactor is adopted from [73, 74], with some modifications. Within the biofilm, the cell growth leads to the production of acetate as a metabolic product. A primary modeling approach in the aforementioned papers assumes the total biomass has a constant concentration and exists in two forms, active and inactive, each of which occupies some volume fraction. We assume that biomass exists only in active form, thus the equations describing the volume-fraction change within the film effectively become a single equation for the rate of change of film thickness, *L*_*f*_, which is a differential state in our system. Moreover, the film growth is affected by a constant detachment rate. It is also assumed that the reaction occurs only within the biofilm, so the only source of acetate in the bulk phase comes from exchange with the biofilm through the boundary layer. We further assume the transport-reaction phenomena in the biofilm are much faster than the transport that occurs across the boundary layer and in the bulk phase; accordingly, the conservation laws inside the biofilm are considered to be in pseudo steady-state [73]. Hence, the computational model consists of a set of nonlinear secondorder ordinary differential equations that describe the spatial distribution of substrate, acetate and overpotential within the biofilm, coupled with a set of first-order ordinary differential equations that describe the concentration of CO_2_ in the bulk phase *S*_*b*_, the acetate concentration in the bulk phase *P*_*b*_, and the biofilm thickness *L*_*f*_. As such, the three state variables of the system are described by

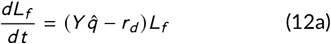

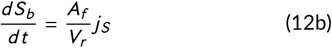

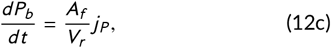

where 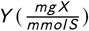 is the biomass yield coefficient, 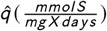 represents an average substrate consumption specific rate within the biofilm, 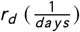 is a detachment rate, *A*_*f*_ (*cm*^2^) is the cross-sectional area of the biofilm, and *V*_*r*_ (*cm*^3^) is the bioreactor volume. The mass balances for the substrate and product are a function of the flux of each species across the linear boundary layer described by

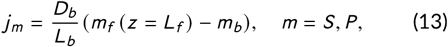

where *m* denotes the species (i.e., substrate and product), 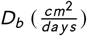 is the diffusivity coefficient in the boundary layer and *L*_*b*_ (*cm*) is the thickness of the boundary layer. The subscript *f* denotes the species concentration in the film at position *z* = *L*_*f*_. The equations that describe the diffusion phenomena within the film are given in the SI. In order to determine the concentrations at *L*_*f*_, a boundary value problem (diffusion within the film) must be solved at each time step, as the concentrations in the biofilm are a function of the bulk concentrations. The computational model is fairly expensive for UQ analyses that rely on Monte Carlo sampling; each model run takes on average 4.5 minutes. The model is subject to time-invariant uncertainty in its parameters and initial conditions. Specifically, the model uncertainty comprises of the conductivity of the biofilm *k*_*bio*_, the maximum growth rate *μ*_*max*_ of the Nerst-Monod model, the yield *Y*, the Monod affinity constant *K*_*s*_, as well as the acetate production-related parameters *α* and *β*. These six uncertain parameters are assumed to follow a uniform probability distribution. Their nominal values are [*k*_*bio*_, *μ*_*max*_, *Y, K*_*s*_, *α, β*]^⊤^ = [1 × 10^−3^, 4.5, 0.25, 3.0, 0.1, 2 × 10^−5^]^⊤^, while they vary uniformly ±10% from the nominal values.

In this case study, we construct data-driven flowmap models of the PCK form in Eq. (5) for the QoIs ***y*** = [*L*_*f*_ *S*_*b*_ *P*_*b*_]^⊤^, such that the six sources of uncertainty constitute the vector of input variables ***x*** in Eq. (5). The three flow-map models, one for each QoI, were trained using simulation data generated via the computational model for lag times in the range of *δ* = [0.05, 0.1] days, which allow us to adequately capture the bioreactor dynamics. Notice that clearly the lag time *δ* must always be larger than the integration time step of the computational model.

The training dataset consists of full state trajectories, as well as one-step ahead samples of the states. We initially generate *N*_*T*_ = 30 trajectories, with fixed uncertain parameters in time, over a process time span from 0 to 3.5 days, which corresponds to approximately *T* = 50 samples per trajectory. Then, using the states ***s***_*k*_ corresponding to each sample ***w***_*k*_, we randomize the uncertain parameters and perform one-step ahead simulations. In this way, approximately 1,400 training samples were generated, while 800 samples are used for training the PCK models. The rationale behind not randomizing the states is that the validation trajectories (step 0 of Figure 2) indicate that there is a high correlation among state values. For instance, as *L*_*f*_ grows in time (under insignificant detachment), *S*_*b*_ decreases due to consumption. Thus, for a given set of uncertain parameters and initial states, a few full state trajectories will help generate more informative training samples. Figure 8 shows the predicted trajectories using the datadriven flow-map PCK model for a given realization of uncertainty and initial conditions, while the true trajectory is juxtaposed. The trajectories correspond to a time-march of 50 steps ahead. We observe a perfect agreement between the predicted and validation trajectories, with the average MTE for the three states being approximately 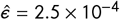.

**FIGURE 8.**
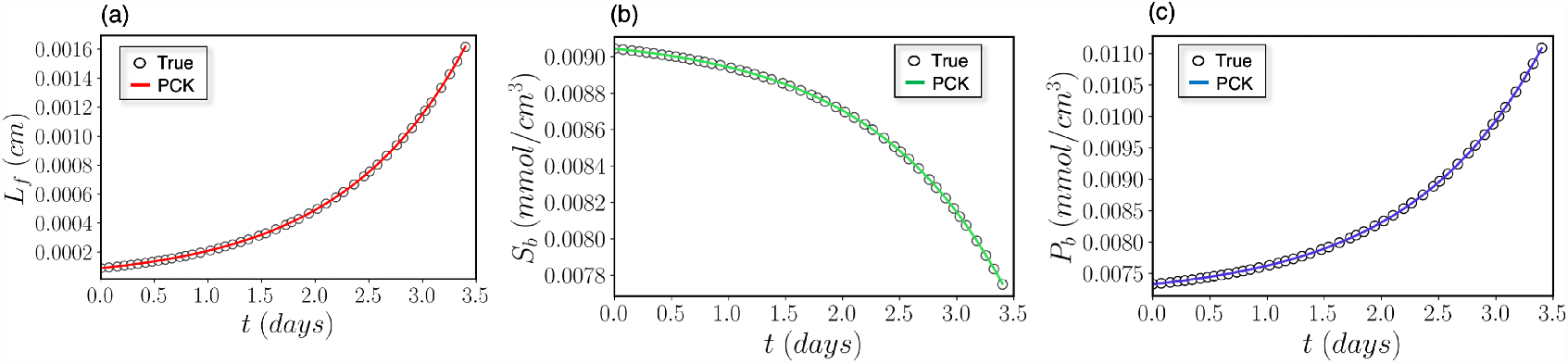
Predicted state trajectories of the the microbial electrosynthesis bioreactor: (a) biofilm thickness, *L*_*f*_, (b) CO_2_ concentration in the bulk phase, *S*_*b*_, and (c) acetate concentration in the bulk phase, *P*_*b*_. Hollow points represent the validation trajectories, while the solid lines represent the trajectories predicted by the PCK-based flow-map models.

An important remark should be made here regarding the benefits of the presented flow-map approach to surrogate modeling of dynamical systems in comparison with the so-called time-frozen approaches discussed in Section 1. First, the flow-map models provide the flexibility to approximate the distribution of states at any time instant of interest without the need for constructing a separate surrogate model for each time instant, as in time-frozen surrogate modeling. For example, if we were to use a time-frozen approach, 50 separate PCK models would need to be constructed for each QoI to predict the time-evolution of the QoI distribution over the 50 time instants considered here. Thus, not only a flow-map modeling approach significantly reduces the number of surrogate models that must be constructed to only one model for each QoI, it also provides flexibility via alleviating the need to build the models at prespecified time points. Furthermore, the flow-map modeling approach enables more efficient data generation. To clarify this point, let us assume that *N*_*p*_ realizations of uncertainty are sufficient for generating a rich training dataset that yields surrogate models with low approximation error. In the case of the time-frozen approach, we would require to generate *N*_*p*_ full state trajectories since the states must be observed at all time instants for all uncertainty realizations. This approach to data generation can become prohibitively expensive, in particular when data generation relies on expensive simulations. However, training the flow-map models, in principle, requires simulation of a limited number of full state trajectories (in this study, 25 trajectories), whereas *N*_*p*_ training samples can be straightforward generated via one-step ahead integration of the computational model. In the following, the use of PCK-based flow-map models is demonstrated for expensive UQ analyzes.

### 4.1 Forward Uncertainty Propagation and Global Sensitivity Analysis

Here, we use the data-driven flow-map models for efficient uncertainty propagation via sample-based approximation of the distribution of the three QoIs. Figure 9(a)-(c) shows the distribution of the QoIs at *t* = 3.5 days. To approximate the distribution of QoIs, the flow-map models were evaluated using 20,000 realizations of the model uncertainty ***x***. Each run of the data-driven flowmap model takes on average less than 3 seconds,^1^ as opposed to the average run time of 4 minutes of the computational model. This implies that the flow-map models significantly accelerate the uncertainty propagation, enabling an approximately 100-fold increase in the computational speed. This is especially beneficial when the distributions are skewed (or bi-modal), as in Figure 9(a)-(c). In this case, a large number of samples, *O* (10^4^ − 10^5^), would typically be required for accurate sampled-based approximation of distribution, or statistical moments of QoIs. Although not shown here, we can efficiently approximate the distribution of QoIs at any time instant using trajectories generated by the surrogate model.

**FIGURE 9.**
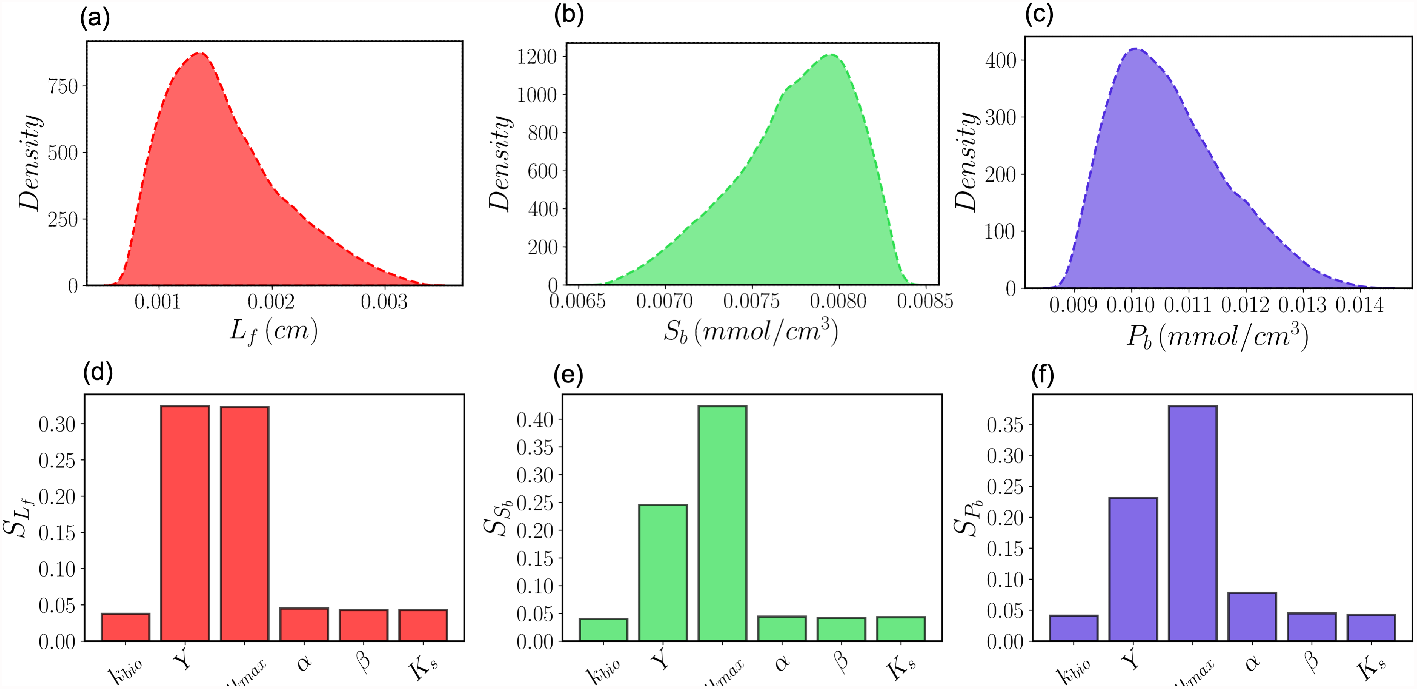
Fast uncertainty propagation and global sensitivity analysis of the the microbial electrosynthesis bioreactor using data-driven flow-map models of quantities of interest. Subplots (a)-(c) show the kernel density estimates of the distribution of the biofilm thickness (*L*_*f*_), concentration of CO_2_ in the bulk phase (*S*_*b*_), and acetate concentration in the bulk phase (*P*_*b*_) predicted by the PCK models at time *t* = 3.5 days. The distributions of *L*_*f*_, *S*_*b*_ and *P*_*b*_ are approximated via Monte Carlo sampling using 20,000 realizations of uncertain model parameters, where a 100-fold computational speedup in sample-based approximation of the distributions is attained. Subplots (d)-(f) show the Borgonovo indices, denoted by S, that quantify the global sensitivity of *L*_*f*_, *S*_*b*_ and *P*_*b*_ at *t* = 3.5 days with respect to the six uncertain model parameters. The Borgonovo indices are approximated based on 20,000 uncertainty realizations.

Moreover, we use the data-driven flow-map models to perform a global sensitivity analysis in order to asses the importance of the six uncertain model parameters, ***x***, on the QoIs ***Y***. This is done via evaluation of the Borgonovo indices [75], denoted by *S*, which are based on the full distribution of QoIs, as opposed to their statistical moments. The results of global sensitivity analysis of QoIs at *t* = 3.5 days are shown in Figure 9(d)-(f), where each bar corresponds to a different uncertain model parameter. The Borgonovo indices are approximated using the same 20,000 samples used in forward UQ. We observe that the probabilistic uncertainty of yield *Y* and maximum growth rate *μ*_*max*_ have the most dominant effects on the variability of the three QoIs, while the product concentration *P*_*b*_ is also significantly affected by the uncertainty in the parameter *α*, which is the metabolism-related productivity constant.

### 4.2 Bayesian Inference of Unknown Model Parameters

We now use the data-driven flow-map models to solve a Bayesian inference problem in order to infer the uncertain model parameters ***x***. Bayesian inference relies on Bayes theorem to estimate the posterior probability distribution of the unknown model parameters from available data. Here, noisy observations of *L*_*f*_, *S*_*b*_ and *P*_*b*_ at time instants {0.5, 1.0, 1.5, 2.0, 2.5, 3.0, 3.5} days constitute the dataset D used for parameter inference; measurement noise is modeled as a Gaussian distribution with zero mean and state-dependent variance. Once a vector of system measurements ***d*** at a time instant is observed, the change in our knowledge about the unknown parameters is described by Bayes’ rule [76]

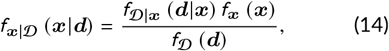

where *f*_***x***|𝒟_ denotes the posterior distribution of the uncertain parameters after observing the data; *f*_𝒟|***x***_ is the likelihood function that describes the probability of observing data given the parameter estimates; *f*_***x***_ is the prior distribution of parameters; and *f* _𝒟_ is the so-called evidence or marginal likelihood that ensures the posterior distribution integrates to 1.

As Eq. (14) implies, Bayesian inference provides an explicit representation of the uncertainty in the parameter estimates via characterizing the full posterior distribution of unknown parameters ***x***. The prior distribution of parameters and the likelihood function must be specified to solve Eq. (14). Here, we used the same uniform distributions as those used to construct the PCK surrogate models to represent the prior distributions, although these can be different. The likelihood function is specified by the observation noise model, which is assumed to be zero-mean Gaussian with state-dependent variance in this work. We use a particle filtering method, namely sequential Monte Carlo (SMC) [77], to approximately solve the Bayesian inference problem by iteratively updating the posterior *f*_***x***|𝒟_ at every time instant that system observations become available; see [43] for further details. Notice that parameter estimation via Bayesian inference methods such as SMC relies on accurate construction of the probability distributions in Eq. (14). As described in Section 4.1, the data-driven flow-map models enable efficient sample-based approximation of the distributions using a very large number of samples, which otherwise could be impractical using an expensive computational model.

Figure 10 shows the posterior distribution of the parameters ***x*** at *t* = 3.5 days estimated via SMC using the dataset 𝒟, as specified above. The posterior distributions are approximated using 20,000 particles. Note that the posterior distribution ranges seem to be larger than the prior in some cases, which is an artifact of the kernel density estimation (i.e., the selection of the bandwidth parameter) [78]. Figure 10 suggests that only the posterior distributions of parameters *Y* and *μ*_*max*_ have changed significantly with respect to their priors. It is also evident that the mean of the posterior distributions (blue vertical lines) for parameters *Y* and *μ*_*max*_ provides a fairly accurate estimate for the true, but unknown, parameter values (brown vertical lines). In particular, the true value and the posterior mean are indistinguishable, while the posteriors are much more narrow compared to priors as stated before. Nonetheless, the posterior distributions for the other parameters remain similar to their priors with little to no change, suggesting these parameters cannot be estimated using the available dataset 𝒟. This can be attributed to the lack of information content of system observations 𝒟 for inferring the unknown parameters; a deficiency that can be addressed via optimal experiment design [79, 80]. We again note the flexibility of the flow-map models that would allow us to seamlessly add new observation points, should that become necessary for better parameter inference, without the need to construct new surrogate models for the QoIs observed at new time points.

**FIGURE 10.**
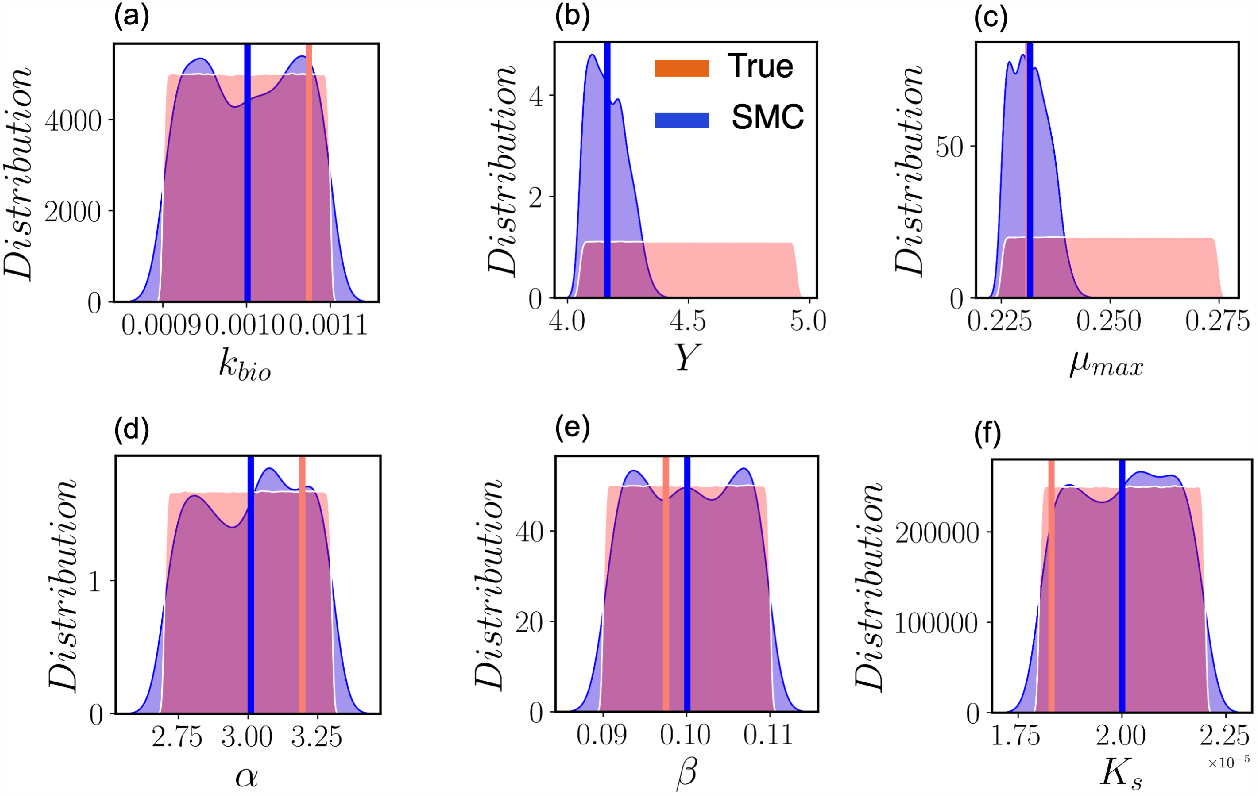
Bayesian inference of unknown parameters of the computational model of the microbial electrosynthesis bioreactor. The parameters are estimated via sequential Monte Carlo using 20,000 particles. Red and blue distributions represent the prior and posterior distributions of the unknown model parameters at time 3.5 days, respectively. The red vertical lines correspond to the true parameters, while the blue vertical lines are the estimated posterior mean value of parameters.

## 5 CONCLUSIONS

This paper presented a flow-map modeling approach based on polynomial chaos Kriging for the discovery of system dynamics from data. Data-driven flow-map models directly approximate the integration operator of differential equations that describe the state transitions of a dynamical system as a function of system state and input variables. We illustrated the usefulness of the proposed approach for learning mathematical descriptions of nonlinear dynamical systems and deriving dynamic surrogate models for fast uncertainty quantification applications. Our analyzes reveal that polynomial chaos Kriging-based flow-maps offer significant benefits in terms of data efficiency, as well as computational efficiency of data generation, for the discovery of nonlinear system dynamics and surrogate modeling.

## Data Availability

All software required for reproducing the case studies presented is available through the CUBES github organization at https://github.com/cubes-space/ DataDriven-FlowMaps and any additional data is available upon request.

## Authorship Contributions

GM and AM conceived the concept of this contribution. GM performed the analysis with help from AJB and FS and oversight from AM, APA, and DSC. GM, AM, and AJB wrote the manuscript. All authors edited the manuscript.

## Acknowledgements

This material is based upon work supported by NASA under grant or cooperative agreement award number NNX17AJ31G.

## 6 SUPPLEMENTARY INFORMATION

### 6.1 Additional Parameters for Morris-Lecar

The extra expressions that appear in the Morris-Lecar model are given by

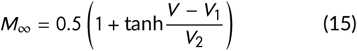

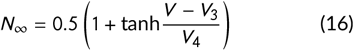

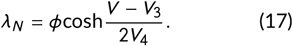

The Type I model parameters are given as

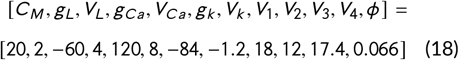

### 6.2 Extra information for the MES Reactor

Here we provide the complete mathematical description of the MES reactor. For completeness we repeat the main equations that were presented in the manuscript. The bulk quantities, with a subscript *b*, are described by:

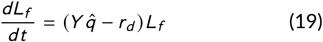

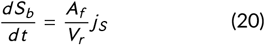

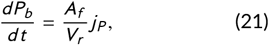

where the various variables are described in the text. The mass balances for the substrate and product are a function of the flux of each species across the linear boundary layer, given by

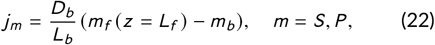

where *m* denotes the species (substrate and product,respectively), *D*_*b*_ is the diffusivity coefficient in the boundary layer and *L*_*b*_ is the thickness of the boundary layer. The subscript *f* denotes the species concentration in the film, at position *z* equal to the film thickness *L*_*f*_.

Within the biofilm, the local growth rate follows a modified Monod-Nerst law given by

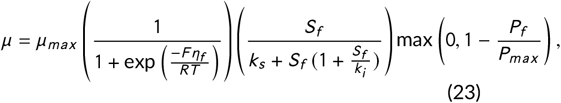

where *k*_*s*_ is the affinity/Monod constant, *k*_*i*_ is a substrate inhibition constant and *P*_*max*_ is a maximum product concentration. *F* is the Faraday constant. Therefore, we consider inhibition both by the substrate and the product. Under the pseudo-steady state assumptions, the following differential equations are solved within the biofilm:

#### Substrate Concentration

We have the differential conservation law, the boundary condition at the biofilm edge as well as a no flux condition at the cathode given by

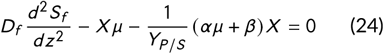

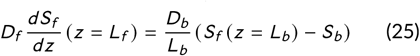

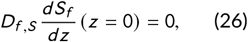

respectively

#### Product Concentration

Similar to the substrate, we have the following equations

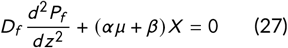

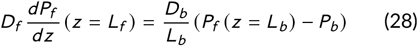

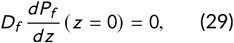

Note that it has been assumed that the diffusivity of substrate and product within the film is equal, denoted by *D*_*f*_.

#### Overpotential

Finally, we have the equations for the overpotential; We describe the diffusion of electrons in the conductive biofilm, we have a fixed voltage at the cathode and no-flux at the biofilm edge

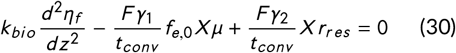

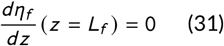

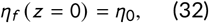

respectively.

## Footnotes

1 Notice that the evaluation time of a PCK model depends on a multitude of factors, such as the degree of the polynomial basis functions, kernel type, and, mainly, amount of data used to train the model. Additionally, a kernel-based model such as PCK is more expensive to evaluate than a polynomial chaos expansion.

